# Heterogeneity in Neuronal Dynamics is Learned by Gradient Descent for Temporal Processing Tasks

**DOI:** 10.1101/2022.01.19.476851

**Authors:** Chloe N. Winston, Dana Mastrovito, Eric Shea-Brown, Stefan Mihalas

**Affiliations:** Department of Neuroscience, University of Washington, Seattle, WA; Department of Computer Science, University of Washington, Seattle, WA; University of Washington Computational Neuroscience Center, Seattle, WA; Allen Institute for Brain Science, Seattle, WA; Department of Applied Mathematics, University of Washington, Seattle, WA

**Keywords:** after-spike currents, generalized-leaky-integrate-and-fire model, neuronal heterogeneity, rate-based neuron model, recurrent neural network

## Abstract

Individual neurons in the brain have complex intrinsic dynamics that are highly diverse. We hypothesize that the complex dynamics produced by networks of complex and heterogeneous neurons may contribute to the brain’s ability to process and respond to temporally complex data. To study the role of complex and heterogeneous neuronal dynamics in network computation, we develop a rate-based neuronal model, the generalized-leaky-integrate-and-firing-rate (GLIFR) model, which is a rate-equivalent of the generalized-leaky-integrate-and-fire model. The GLIFR model has multiple dynamical mechanisms which add to the complexity of its activity while maintaining differentiability. We focus on the role of after-spike currents, currents induced or modulated by neuronal spikes, in producing rich temporal dynamics. We use machine learning techniques to learn both synaptic weights and parameters underlying intrinsic dynamics to solve temporal tasks. The GLIFR model allows us to use standard gradient descent techniques rather than surrogate gradient descent, which has been utilized in spiking neural networks. After establishing the ability to optimize parameters using gradient descent in single neurons, we ask how networks of GLIFR neurons learn and perform on temporally challenging tasks, such as sinusoidal pattern generation and sequential MNIST. We find that these networks learn a diversity of parameters, which gives rise to diversity in neuronal dynamics. We also observe that training networks on the sequential MNIST task leads to formation of cell classes based on the clustering of neuronal parameters. GLIFR networks have mixed performance when compared to vanilla recurrent neural networks but appear to be more robust to random silencing. When we explore these performance gains further, we find that both the ability to learn heterogeneity and the presence of after-spike currents contribute. Our work both demonstrates the computational robustness of neuronal complexity and diversity in networks and demonstrates a feasible method of training such models using exact gradients.

## 1 Introduction

Artificial neural networks (ANNs) have been used to emulate the function of biological networks at a system level (Yamins et al., 2014; Rajan, Harvey, & Tank, 2016). Such models rely on the fantastic capacity of ANNs to be trained to solve a task via back-propagation, using tools developed by the machine learning community (Goodfellow, Bengio, & Courville, 2016). However, the way in which the neurons in recurrent ANNs integrate their inputs over time differs from neuronal computation in the brain (Figure 1A-B). Primarily, biological neurons are dynamic, constantly modulating internal states in a nonlinear way while responding to inputs. A biological neuron maintains a membrane potential that varies not only with input currents but also through history-dependent transformations. When its voltage exceeds some threshold, a neuron produces a spike, a rapid fluctuation in voltage that is considered the basis of neuronal communication. While these are the defining features of a neuron, neurons exhibit additional, more complex types of dynamics, including threshold variability and bursting. Threshold adaptation gives rise to threshold variability and thus modulates the sensitivity of a neuron to inputs. A proposed mechanism is that the threshold fluctuates in a voltage-dependent manner, possibly due to the gating mechanisms of various ion channels (Fontaine, Peña, & Brette, 2014). Another type of dynamic is responsible for bursting and related spiking behaviors. Bursting results from depolarizing currents which can be induced by prior spikes. A broader array of spiking patterns can be explained by considering both hyperpolarizing and depolarizing after-spike currents, currents that are modulated by a neuron’s spiking activity (Mihalaş & Niebur, 2009).

**Figure 1:**
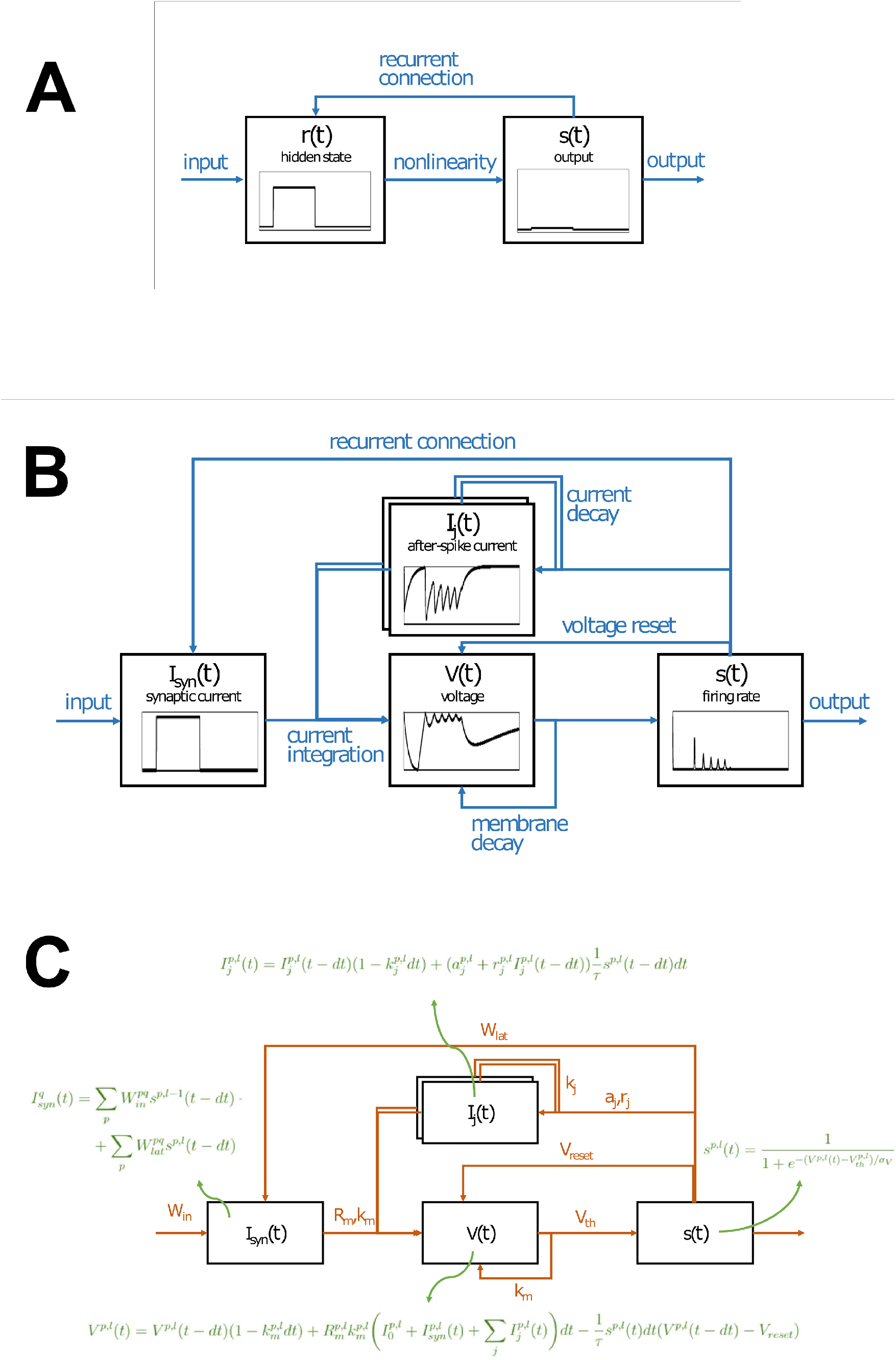
A. Schematic of a “vanilla” RNN cell or neuron. A RNN neuron maintains a hidden state *r*(*t*) that is computed at each timestep by linearly weighting the input signal and the previous output of itself and neighboring neurons through a recurrent connection. The output *s*(*t*) is computed by applying a nonlinear transformation (e.g., tanh or sigmoid) to *r*(*t*). B-C. Schematics of GLIFR neuron. Each neuron maintains a synaptic current *I*_*syn*_ that is computed at each timestep by linearly weighting (*W*_*in*_) the input signal and the previous output of itself and neighboring neurons through a recurrent connection (*W*_*lat*_). The neuron’s voltage *V* decays over time according to membrane decay factor *k*_*m*_ and integrates synaptic currents and after-spike currents *I*_*j*_ over time based on membrane resistance *R*_*m*_ and membrane decay factor *k*_*m*_. Additionally, the voltage tends towards *V*_*reset*_ through a continuous reset mechanism based on the firing rate at a given time. An exponential transformation of the difference between the voltage and the threshold voltage yields a continuous-valued normalized firing rate, which varies between 0 and 1. The normalized firing rate, along with terms *a* _*j*_ and *r*_*j*_, is used to modulate the after-spike currents that decay according to decay factor *k*_*j*_. The dynamics present in a GLIFR neuron give rise to its key differences from RNN neurons; GLIFR neurons can express heterogeneous dynamics, in contrast to the fixed static transformations utilized in RNN neurons.

In sum, biological neuronal dynamics can be thought of as continuous, nonlinear transformations of internal neural states, such as ionic currents and voltage. This is in contrast to a typical artificial neuron that maintains a single state resulting from a static, linear transformation of an input vector. Moreover, diversity in intrinsic neuronal mechanisms give rise to heterogeneity in the dynamics that biological neurons express. Such heterogeneity is generally absent in artificial neural networks. For example, not all neurons in the brain display bursting behavior or threshold adaptation. The brain also exhibits diversity in neuronal properties, such as ionic conductances, threshold, and membrane properties. This diversity is evident from experiments fitting neuronal models to spiking behavior observed in the mouse cortex (Teeter et al., 2018). Heterogeneity of such neural characteristics further amplifies the diversity in neuronal dynamics. In contrast, the only diversity in typical ANNs lies in synaptic weights. Typical recurrent neural network (RNN) neurons do not exhibit intrinsic dynamics, and each neuron responds identically to input with a common, fixed activation function.

Yet, vanilla RNNs which employ linear weighting of inputs and nonlinear transformation of hidden states over time (Figure 1A) have managed to perform quite well on complex tasks without incorporating the complexity and heterogeneity prominent in biological networks. For example, variations of RNNs have been successful at classifying text (Liu, Qiu, & Huang, 2016) and classifying images from a pixel-by-pixel presentation (Goodfellow et al., 2016; Li, Li, Cook, Zhu, & Gao, 2018; LeCun, Bengio, & Hinton, 2015). This raises the question, what, if any, is the advantage of neuronal dynamics and the diversity thereof? Do neuronal dynamics enable the modeling of more temporally complex patterns? Does neuronal heterogeneity improve the learning capacity of neural networks?

We propose that both complex neuronal dynamics, such as after-spike currents, and heterogeneity in neuronal dynamics across a network would enhance a network’s capacity to model temporally complex patterns. Moreover, we hypothesize that heterogeneity in dynamics will be optimal and will be learned when parameters are optimized in addition to synaptic weights during training. In order to evaluate these hypotheses, we develop and test a neuronal model whose parameters can be optimized through gradient descent.

A set of approaches has been previously used to develop neuronal models which encapsulate the above-mentioned biological complexities. These range from models that include a description of channels (Hodgkin & Huxley, 1952; Morris & Lecar, 1981) to more compact models that synthesize the complexities into more abstract dynamical systems (Izhikevich, 2003) or hybrid systems (Mihalaş & Niebur, 2009). Networks of such neurons have been constructed (Markram et al., 2015; Billeh et al., 2020), but they are difficult to optimize, either to solve a task or to fit biological data.

Small increases in dynamical complexity can have significant benefits. Adaptation dynamics can help neural networks perform better on predictive learning tasks (Burnham, Shea-Brown, & Mihalas, 2021) and achieve better fits of neuronal and behavioral data in mice (Hu et al., 2021). Inroads have been made to allow the optimization of spiking models using backpropagation via surrogate gradients (Huh & Sejnowski, 2018; Neftci, Mostafa, & Zenke, 2019). Such methods have revealed the importance of neuronal integration and synaptic time scales (Perez-Nieves, Leung, Dragotti, & Goodman, 2021) and adaptation (Salaj et al., 2021; Bellec et al., 2020) in network computation. Previous models (Perez-Nieves et al., 2021; Burnham et al., 2021) have also shown the importance of time-scale diversity in temporal tasks. However, these models are still significantly simpler than those found to fit single neuron data well (Teeter et al., 2018). To allow larger scale dynamics to be fit well, we want to be able to optimize neural networks with the type of dynamics proven to fit such complexities. Here we focus on the addition of after-spike currents, which can lead to a wide set of observed behaviors, including bursting (Gerstner, Kistler, Naud, & Paninski, 2014; Mihalaş & Niebur, 2009).

While spiking neurons are more biologically realistic, they are generally more difficult to optimize. Thus, we develop a model which is typically differentiable, but becomes a spiking model in the limit of taking a parameter to 0. We refer to this novel neuronal model as the generalized leaky-integrate-and-firing-rate (GLIFR) model. The GLIFR model is built on the spike-based generalized-leaky-integrate-and-fire (GLIFR) model (Teeter et al., 2018) and produces the equivalent of after-spike currents, dynamics dependant on a neuron’s firing history. Unlike the GLIF model, the differentiability of the GLIFR model enables the application of standard deep learning techniques to optimize parameters underlying intrinsic neuronal dynamics. We use gradient descent to optimize networks of GLIFR neurons on several temporal tasks and assess the performance and robustness of these networks. We find that it is possible to optimize both intrinsic neuronal parameters and synaptic weights using gradient descent. Optimization of neuronal parameters generally leads to diversity in parameters and dynamics across networks. Moreover, when we test several variations of the GLIFR model, we find that both the presence of neuronal dynamics and the heterogeneity in neuronal properties improve performance. While the GLIFR networks are outperformed by long-short-term-memory (LSTM) networks, our networks have mixed performance when compared to vanilla RNNs on a pattern generation task and a temporally complex sequential MNIST task. In sum, we develop a method to optimize neuronal parameters in individual neurons and thereby suggest a computational role of neuronal complexity and heterogeneity in the brain. We provide code for creating and optimizing GLIFR models in Python.

## 2 Related Work

In our work, we assess how, if at all, the presence of neuronal dynamics, as well as the heterogeneity thereof, confers performance improvements in RNNs. Prior work has developed RNNs that express biologically realistic dynamics, giving rise to a class of networks known as spiking neural networks (SNNs). One example of an SNN (Zenke & Vogels, 2021) uses a leaky-integrate-and-fire dynamic across its neurons. In these neurons, the membrane potential was modulated according to Equation 1 where *S* represents whether the neuron is spiking, and *V* represents the neuron’s membrane potential.

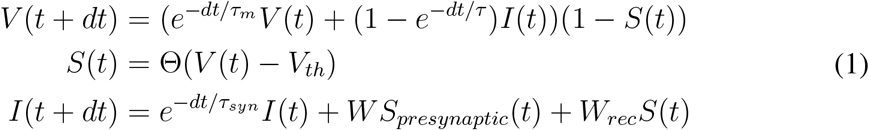

At each timestep, the voltage exponentially decays towards 0 while being increased by the neurons’ synaptic currents that also decay exponentially. Spiking drives the voltage to 0. Only synaptic weights were trained, and a surrogate gradient was employed to address the difficulty presented by the undifferentiable Heaviside function. Specifically, when computing gradients of the loss with respect to parameters, the gradient of the Heaviside function was approximated by a smoother function (Equation 2). Modifying the gradient descent technique only to allow for this surrogate gradient, (Zenke & Vogels, 2021) achieved high performance on an MNIST task.

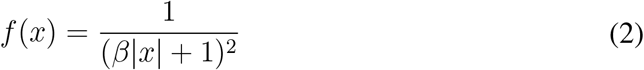

Further work has studied the effect of spike frequency adaptation on network performance, using models of after-spike currents. Spike-frequency adaptation improved the performance of LIF SNNs on temporally challenging tasks, such as a sequential MNIST task where the network must classify images of handwritten images based on a pixel-by-pixel scan, an audio classification task where the network must identify silence or spoken words from the Google Speech Commands Dataset, and an XOR task where the network must provide the answer after a delay following the inputs (Salaj et al., 2021). These networks were found to approach RNN performance on the second audio classification task.

The work described thus far used spiking models of biological dynamics and did not explore the advantage heterogeneity could confer. Several approaches have been taken to this end. One approach is to initialize an SNN with heterogeneous dynamics but optimize only synaptic weights. This method achieved comparable or higher performance than ANNs that employed convolutional and/or recurrent transformations on an object detection task (She, Dash, Kim, & Mukhopadhyay, 2021). A second approach is to optimize intrinsic neuronal parameters in addition to synaptic weights. Under the hypothesis that neuronal heterogeneity is computationally advantageous, the learned parameters will be diverse across the trained network. To this end, one study extended the surrogate gradient technique for LIF SNNs to also optimize membrane and synaptic time constants across networks (Perez-Nieves et al., 2021). It was found that these networks learned heterogeneous parameters across the network when trained on temporal MNIST tasks, a gesture classification task, and an auditory classification task. On some tasks, learning parameters improved performance over learning synaptic weights alone. Learning parameters relied on surrogate gradients. Our work establishes a method of training neuronal parameters with standard gradient descent by using a rate-based neuronal model; moreover, this model additionally expresses after-spike currents as schematized in Figure 1A-B.

To summarize, previous work has suggested that heterogeneity in integration and synaptic time scales improves network performance. This opens the door to the questions we address here: whether after-spike currents that produce complex dynamics within individual neurons can be trained, whether these after-spike currents improve network performance, and whether training naturally leads to their heterogeneity. It also invites the question of whether neuronal models with these complex dymamics can be designed that learn without employing a surrogate gradient technique. In what follows, we show that this is possible, and illustrate how this results in heterogeneous neuronal dynamics as well as mixed performance relative to vanilla RNNs.

## 3 Model for After-Spike Currents

We develop a neuronal model that exhibits after-spike currents and is conducive to gradient descent (Figure 1B, C). To do this, we build on a previously described spike-based GLIF model that incorporates after-spike currents (Teeter et al., 2018). Although prior work has used surrogate gradient methods for optimizing discrete spiking neural networks, we transform this GLIF model into what we term a GLIFR model, which is rate-based to facilitate optimizing neuronal parameters with traditional gradient descent mechanisms. Specifically, we modify the GLIF_3_ model from (Teeter et al., 2018) to produce firing rates (Equation 4) rather than discrete spikes (Equation 3). This rate-based approach enables the use of exact gradients rather than surrogate gradients across the spiking function. Additionally, it is more akin to vanilla RNNs that utilize smooth activation functions, thereby motivating the application of standard deep learning techniques.

Below, we describe how state variables are computed at each timestep in the GLIFR model, comparing the computation to that in the GLIF model. We use a subscript *s* for the GLIF state variables and no subscript for the equivalent GLIFR variables. We use superscript *p* to represent the index of the neuron in consideration and *l* for the associated layer where postsynaptic layers are indexed with larger numbers than are presynaptic layers. For example, *s*^*p,l*^(*t*) defines the normalized firing rate of the *p*^*th*^ neuron in the *l*^*th*^ layer.

The discrete spiking equation can be expressed as in Equation 3. H refers to the Heaviside function where ℍ(*x*) = 1 when *x >* 0 and ℍ(*x*) = 0 when *x≤* 0. In the GLIFR model, we define *s*^*p,l*^(*t*), the normalized firing rate that is unitless and varies between 0 and 1 (Equation 4). A raw firing rate 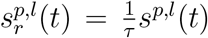 may be derived where *τ* (ms) represents how often we check if the neurons has crossed the threshold, and we set *τ* to *dt* throughout the study. While *s* represents a normalized firing rate, for simplicity of terminology we will refer to it as the firing rate. The parameter *σ*_*V*_ (mV) controls the smoothness of the voltage-spiking relationship, with low values (*σ*_*V*_ *<<* 1*mV*) enabling production of nearly discrete spikes and higher values enabling more continuous output (Figure 2A, B). We used *σ*_*V*_ = 1*mV* for all simulations, unless otherwise noted. Thus, for the GLIF and GLIFR models, respectively, spiking is defined as follows:

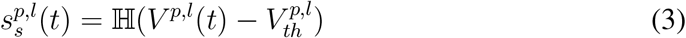

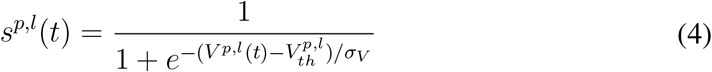

where (3) corresponds to the GLIF and (4) to the GLIFR model.

**Figure 2:**
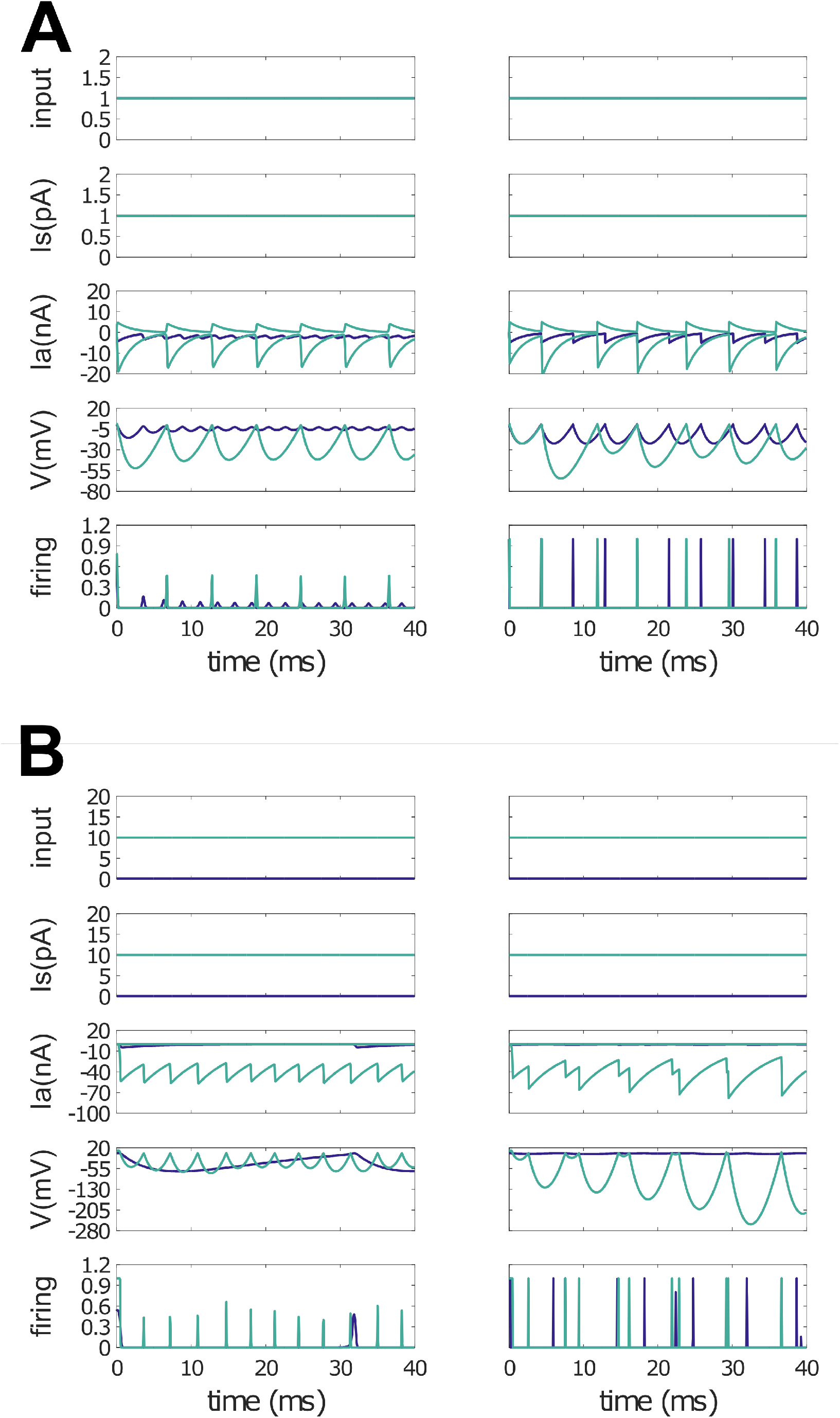
A. Sample responses to constant input. These plots show example outputs of two neurons when provided with a constant input over a 40 ms time period. The two neurons share identical membrane-related parameters and decay factors for after-spike currents. The neuron whose output is traced in purple has the following after-spike current related parameters: *r*_1_, *r*_2_ = −1; *a*_1_, *a*_2_ = − 5000*pA*. The neuron whose output is represented in green has the following parameters: *r*_1_ = − 1, *r*_2_ = 1, *a*_1_ = 5000*pA, a*_2_ = − 5000*pA*. The difference in after-spike current parameters gives rise to different types of dynamics. The lefthand column contains outputs produced by neurons with *σ*_*V*_ = 1, whereas the righthand column contains outputs produced by neurons with *σ*_*V*_ = 0.001*mV*, demonstrating the ability of the GLIFR neuron to produce spike-like behavior while maintaining its general differentiability. B. Sample responses to different amplitude inputs. These plots show example outputs of a neuron when provided with different magnitudes of constant input over a 40 ms time period. Larger inputs appear to yield higher-frequency oscillations in the neuron’s output. The lefthand column contains outputs produced by neurons with *σ*_*V*_ = 1*mV*, whereas the righthand column contains outputs produced by neurons with *σ*_*V*_ = 0.001*mV*.

After-spike currents are modeled as a set of separate ionic currents (corresponding to *j* = 1, 2, …) as in Equation 6. We use two after-spike currents with arbitrary ionic current equivalents, but this model can theoretically be extended to more currents. As in the GLIF model (Equation 5), decay factor 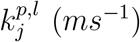 is used to capture the continuous decay of the current, and the “spike-dependent current” is determined by multiplicative parameter 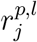 and additive parameter 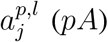. In the GLIFR model though, this “spike-dependent current” is scaled by the raw firing rate 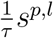.

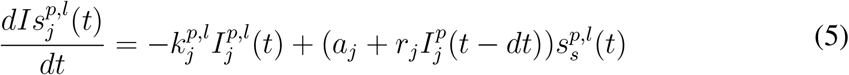

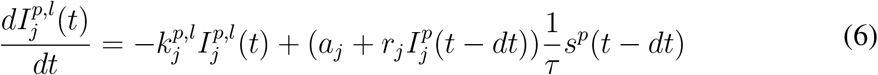

Synaptic currents, currents resulting from input from the previous layer as well as lateral connections, are modeled according to Equation 8. We do not include an exponential decay term for simplicity as we focus on cellular properties, and the model is equivalent in the discrete and continuous space. The first term in this equation represents the integration of firing inputs from the presynaptic layer and the second term describes the integration of firing inputs from the same layer through lateral connections. The latter connection is associated with a synaptic delay denoted Δ*t*. We use Δ*t* = 1*ms*.

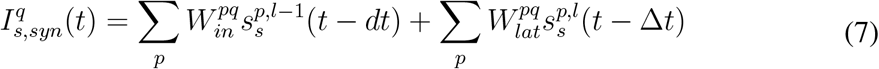

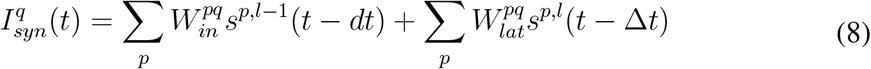

Here, *W*_*pq*_ (pA) defines how much the presynaptic spiking outputs (of neuron *p*) affects the postsynaptic current (of neuron *q*).

Finally, voltage is modeled similarly to in a traditional GLIF model (Equation 9), but in order to preserve the simplicity of our model to facilitate gradient descent, we ignore the more complex reset rule that is used in the GLIF model (Teeter et al., 2018). Instead, we set *f*_*r*_ = 1 in Equation 10.

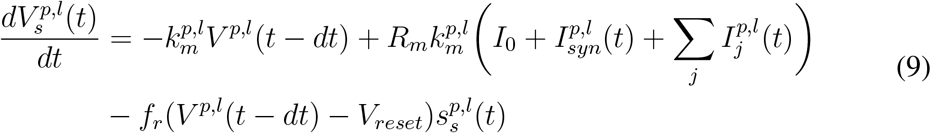

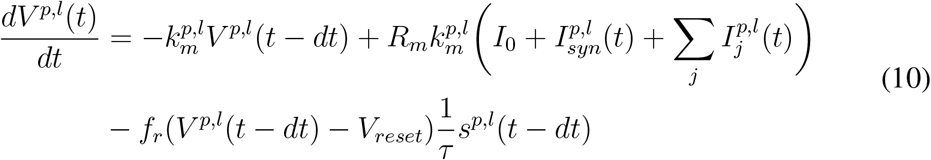

Here, *V* ^*p,l*^(*t*) decays according to *k*_*m*_ (*ms*^−1^). It integrates current based on resistance *R*_*m*_ (*G*Ω). Voltage also tends towards the reset voltage *V*_*reset*_ (*mV*) at a rate proportional to the firing rate *s*^*p,l*^. This is a continuous equivalent of the discrete voltage reset.

Varying the values of 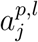 and 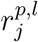 can give rise to a variety of complex patterns including bursting. As shown in Figure 2A, hyperpolarizing values enable the neuronal firing rate to oscillate slightly, and a combination of hyperpolarizing and depolarizing after-spike currents enable regular oscillations in firing rate. Because we model firing rates rather than individual spikes, we take this as a form of bursting. We furthermore find that for a given set of neuronal parameters, larger inputs yield higher-frequency oscillations in firing rate (Figure 2B). We note that in these simulations and later in trained networks, the GLIFR model is theoretically capable of producing both biologically plausible patterns and less biologically realistic activity.

## 4 Method for Training Weights and Parameters in GLIFR Networks

We use standard gradient descent to optimize networks of GLIFR neurons, training both synaptic weights and parameters in each neuron separately, similar to (Perez-Nieves et al., 2021). In this way, the network can learn heterogeneous parameters across neurons. To enable simulating these networks during training, we discretize the after-spike current and voltage equations as shown in Equations 11 and 12 respectively.

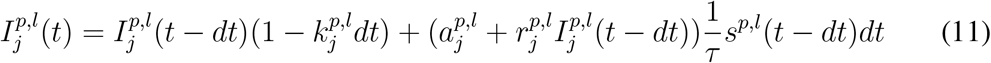

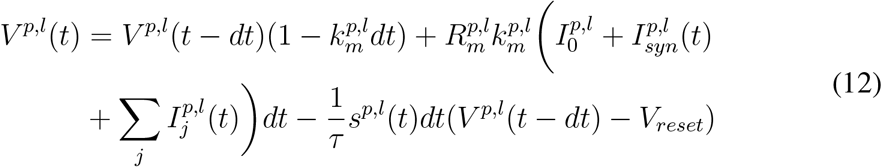

We introduce several changes in variables to enable training multiplicative terms and to promote stability. Because multiplicative terms can pose challenges for optimization by gradient descent, we keep resistance fixed and do not optimize it. Instead, while we do optimize *k*_*m*_, we optimize the product *W*_*pq*_*R*_*q*_*k*_*q*_*dt* rather than *W*_*pq*_ (either for incoming weight or lateral weights) or *R*_*q*_ individually, and we refer to this product as *ω*_*pq*_. This combines the synaptic current and voltage computations.

We also add biological constraints to several parameters to maintain stability through training. In biological systems, *k*_*m*_ and *k*_*j*_ are restricted to be positive since they represent rates of decay. Moreover, to maintain stability, decay factors should be kept between 0 and 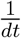 throughout training. To achieve this constraint without introducing discontinuities in the loss function, we optimize 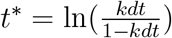 for each decay factor *k*. Thus, *t*\* can be any real number, whereas 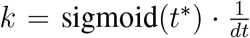 is the decay factor we use when simulating neural activity. In this way, the value of *k* that is used is always between 0 and 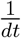.

The term *r*_*j*_ can also introduce instability, so we use a similar method to constrain *r*_*j*_ to [−1, 1]. We optimize 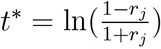 and retrieve *r*_*j*_ using *r*_*j*_ = 1 − 2 *·* sigmoid(*t*\*), which is bounded between −1 and 1.

The units of all states maintained by the GLIFR neuron are shown in Table 1, and the units of all trainable parameters are shown in Table 2. Note that we use *j* = 1, 2 in all our simulations to model two after-spike currents.

**Table 1:** List of states in GLIFR neurons, along with their units.

|  |  |  |
| --- | --- | --- |
| $V(t)$ | voltage | $mV$ |
| $s(t)$ | normalized firing rate | 1 |
| $I_j(t)$ | after-spike current | $pA$ |
| $I_{syn}(t)$ | synaptic current | $pA$ |

**Table 2:** Trained Parameters. List of trainable parameters in GLIFR networks, along with their units. Here, indices *p, l* refer to neuron *p* in layer *l*, so that individual neuron parameters are trained separately.

|  |  |  |
| --- | --- | --- |
| $V_{th}^{p,l}$ | threshold | $mV$ |
| $\omega_{in}^{pq}, \omega_{lat}^{pq}$ | combined synaptic weights | $mV$ |
| $a_j^{p,l}$ | additive constant in after-spike current | $pA$ |
| $r_j^{p,l}$ | multiplicative constant in after-spike current | unitless |
| $k_j^{p,l}$ | decay factor for after-spike current | $ms^{-1}$ |
| $k_m^{p,l}$ | voltage decay factor | $ms^{-1}$ |

We employ a deterministic initialization scheme to decrease the likelihood that the network will learn parameters underlying exploding or saturating firing rates; specifically, we initialize the network with small decay constants and thresholds close to the reset voltage of 0*mV*. We use two initialization schemes in our experiments to test the effects of heterogeneous and homogeneous initialization. Parameters associated with both schemes are described in Table 3. Note that the neuron described is in a layer with *N* neurons.

**Table 3:**
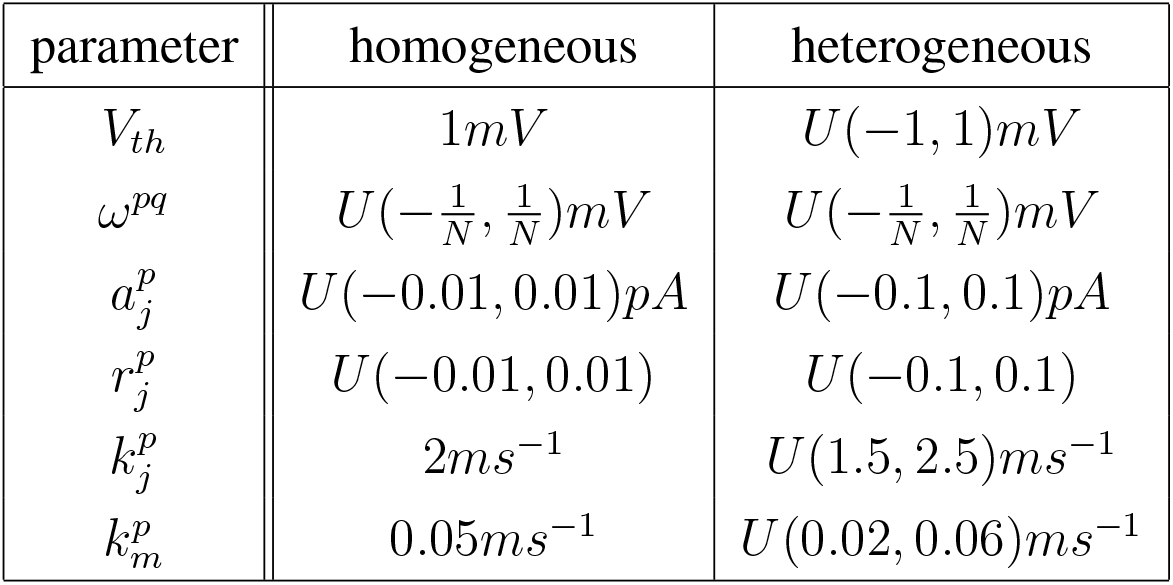
Parameter Initialization. This table describes how each trainable parameter was initialized in GLIFR neurons either homogeneously or heterogeneously across neurons. Note that in both schemes, *ω, r*_*j*_, and *k*_*j*_ were initialized with at least some amount of heterogeneity.

| parameter | homogeneous | heterogeneous |
| --- | --- | --- |
| $V_{th}$ | $1mV$ | $U(-1, 1)mV$ |
| $\omega^{pq}$ | $U(-\frac{1}{N}, \frac{1}{N})mV$ | $U(-\frac{1}{N}, \frac{1}{N})mV$ |
| $a_j^p$ | $U(-0.01, 0.01)pA$ | $U(-0.1, 0.1)pA$ |
| $r_j^p$ | $U(-0.01, 0.01)$ | $U(-0.1, 0.1)$ |
| $k_j^p$ | $2ms^{-1}$ | $U(1.5, 2.5)ms^{-1}$ |
| $k_m^p$ | $0.05ms^{-1}$ | $U(0.02, 0.06)ms^{-1}$ |

In contrast to previous work (Perez-Nieves et al., 2021) with spiking models, we find that standard gradient descent without surrogate gradients suffices to train GLIFR networks. While this relies on the use of a rate-based model rather than a spiking model, GLIFR neurons are capable of producing nearly discrete outputs when *σ*_*V*_ is taken in the limit to 0. Importantly, by utilizing this scheme, we are able to extend our model to account for after-spike currents, a significant, albeit complex, dynamic in biological networks.

## 5 Testing the Optimization of Neuronal Parameters for Learning Realizable Signals

We first confirm the ability of neuronal parameters to be optimized through gradient descent in a theoretically simple task - to learn a target that is known to be realizable by a GLIFR.

As shown in Figure 3A, we initialize a single GLIFR neuron and record its response to a constant input stimulus over a fixed period of time (10 ms with timestep duration of 0.05 ms.) This is our target neuron. We then create a second neuron (learning neuron) that is equivalent to the target neuron only in the incoming weights used. We train the learning neuron to produce the target neuron’s output, allowing it to learn the parameters that were altered. We used Adam optimization with a learning rate of 0.001.

**Figure 3:**
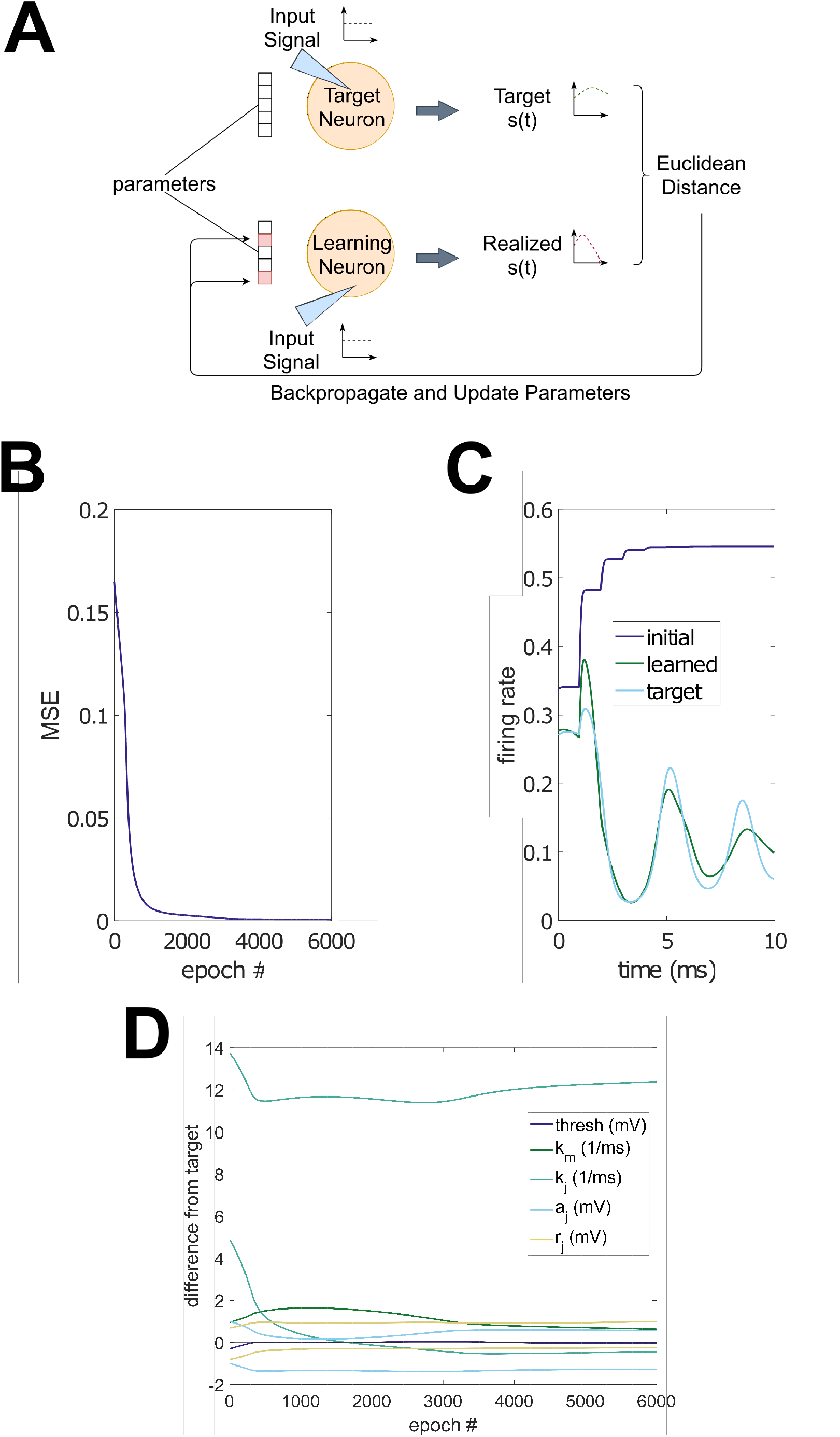
A. Testing optimization in single neurons for realizable signals. In the realizable pattern generation task depicted here, a neuron (the target neuron) is used to create the target firing pattern. A second neuron (the learning neuron) is initialized with different parameters and learns parameters to produce the target. B. Training loss. This plot shows an example trace of mean-squared error over training of a single neuron. C. Output of learning neuron. This plot shows the output of the learning neuron prior to training and after training, along with the target, demonstrating the ability of a single GLIFR neuron to learn simple patterns. D. Learned parameters. This plot depicts the difference between the values of the trainable parameters in the learning neuron and the corresponding values in the target neuron over training epochs. Note that a value of 0 represents equivalence to the target network in a particular parameter.

We tested the ability of the learning neuron to learn the pattern produced by the target neuron. Figure 3B-C show that a single neuron successfully learned to reproduce the dynamics of the target neurons. However, the final parameters learned by the learning neuron differed from the parameters in the target neuron (Figure 3D), illustrating that different internal mechanisms can lead to similar dynamics (Prinz, Bucher, & Marder, 2004). This supports the idea that varying distributions of parameters across a network may nevertheless lead to similar network function.

## 6 Strategy for Analyzing Task Learning and Performance

After verifying the theoretical ability to optimize neuronal parameters in simple GLIFR systems, we turn to more complex tasks. As in previous work (Perez-Nieves et al., 2021), we aimed to assess the role of several factors in network computation while evaluating our networks: (1) the presence of biologically realistic dynamics (i.e., membrane dynamics, voltage reset), (2) the presence of after-spike currents, (3) random heterogeneity of neuronal parameters across the network, (4) the learning of heterogeneity of neuronal parameters across the network. Thus, we utilize multiple network types. First, as a baseline, we use a vanilla recurrent neural network (RNN). The neurons used by RNNs are modeled as in Equation 13. Here, *W*_*ih*_ represents the weights applied to the input signal, *W*_*hh*_ refers to the recurrent weights, and *b* refers to the bias term. In contrast to GLIFR neurons, we set the synaptic delay Δ*t* to *dt* in RNNs in order to enable to modeling of nonlinear transformations at the beginning of a simulation. This enables the RNN to perform better than it would if we used Δ*t* = 1*ms* on all tasks we study.

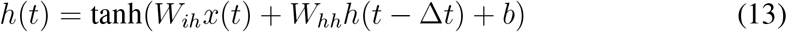

We also use the following variations of the GLIFR network: a GLIFR network with fixed heterogeneity with (FHetA) or without (FHet) after-spike currents, a GLIFR network with refined heterogeneity with (RHetA) or without (RHet) after-spike currents, and a GLIFR network with learned heterogeneity with (LHetA) or without (LHet) after-spike currents. We define fixed heterogeneity as heterogeneity that a network is initialized with and does not alter over training, refined heterogeneity as heterogeneity that a network is initialized with but is altered (‘fine-tuned’) over training, and learned heterogeneity as heterogeneity that a network is not initialized with but is learned over training. These distinctions are illustrated in Figure 4. We finally use an LSTM network as another baseline whose mechanisms improve the ability to model complex temporal dynamics without emulating biology. For each experiment, we set the number of neurons for each network type to yield comparable numbers of learnable parameters.

**Figure 4:**
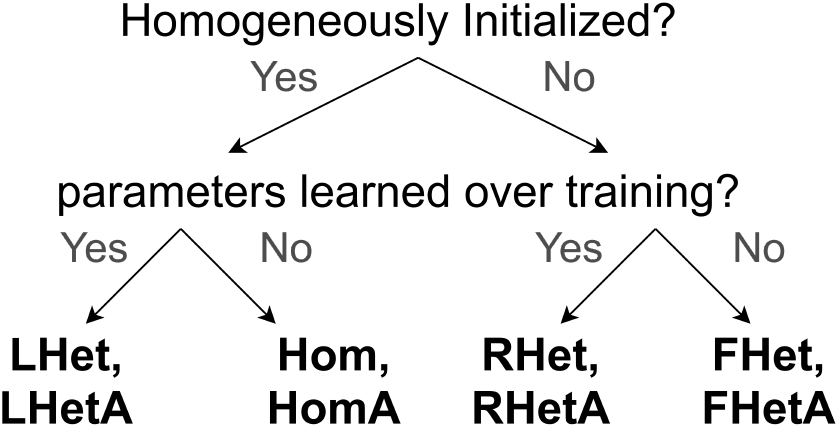
GLIFR schemes. A flowchart describes the variations of the GLIFR networks we explored. These are based on whether the network neuronal parameters were homogeneously initialized or heterogeneously initialized as well as whether these intrinsic neuronal parameters were learned over training. This classification enables us to isolate effects of the expression of after-spike currents, a complex type of dynamic, and the learning of heterogeneous parameters.

Each network, including RNNs, GLIFR networks, and LSTM networks, consists of a single recurrent hidden layer of the appropriate neeuron type, whose outputs at each timepoint are passed through a fully connected linear transformation to yield the final output. The recurrence is incorporated into the GLIFR networks through the term 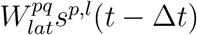 in the synaptic currents equation..

Code for the GLIFR model is publicly available at https://github.com/AllenInstitute/GLIFS_ASC. The model is implemented in PyTorch and thus can be optimized and evaluated like other PyTorch neural network modules.

## 7 Performance on Pattern Generation Task

In order to study the contribution of neuronal complexity and heterogeneity to the ability to learn complex temporal dynamics, we trained networks on a pattern generation task (Figure 5A). Each network consisted of a single recurrent layer of GLIFR, RNN, or LSTM neurons whose outputs were weighted at each timestep to obtain an 1-dimensional time series. The number of parameters and the number of neurons in the hidden layer of each network trained on this task are listed in Figure 5B.

**Figure 5:**
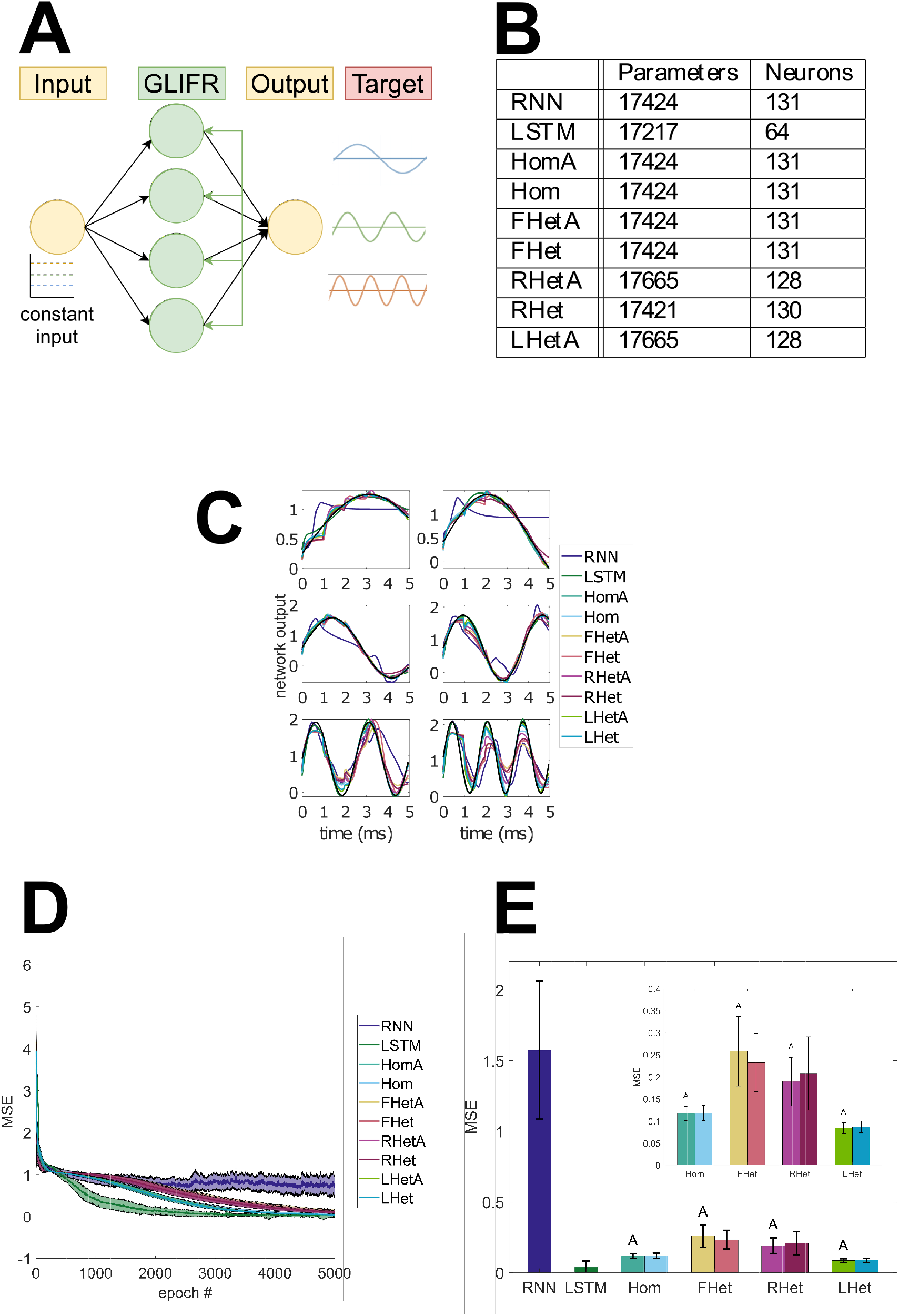
A. Pattern generation task. A schematic shows the pattern generation task. The yellow circles represent signals that are either weighted to form inputs to neurons or result from applying weights to neuronal outputs. The green circles represent neurons. The network is given constant input and is expected to produce a sinusoid whose frequency is directly proportional to the magnitude of the input. B. Metadata for trained networks. The number of trainable parameters is shown for each network along with the size of the hidden layer. C. Sample approximations. These plots depict the target sinusoids in black and the corresponding predictions of sample trained networks. All network types learned the patterns reasonably well. D. Training loss. The mean-squared error of the network on the dataset averaged over thirty random initializations is plotted over training epochs. The shading indicates a moving average of the standard deviation. On average, all network types converged on a solution within the training time. E. Trained model performance. The mean-squared error of each trained network on the pattern generation task is shown. Bars are labeled with RNN, LSTM, Hom, FHet, RHet, or LHet on the x-axis, and an “A” above the bar represents the presence of after-spike currents. For example, the Hom bar with an “A” above it represents the HomA network. This chart shows that the GLIFR networks outperfrom the RNNs but not the LSTM networks. Additionally, learning heterogeneity improved performance of the GLIFR networks, although expression of after-spike currents did not appear to confer performance improvements.

Each network was trained to generate 5 ms sinusoidal patterns whose frequencies are determined by the amplitudes of the corresponding constant inputs. We trained each network to produce six sinusoids with frequencies ranging from 80 Hz to 600 Hz. The *i*^*th*^ sinusoid was associated with a constant input of amplitude (*i/*6) + 0.25. Previous work (Sussillo & Barak, 2013) tackles a more complex version of this task using RNNs but uses Hessian optimization, while we limit our training technique on both the RNN and the GLIFR networks to simple gradient descent. We use an Adam optimizer with a learning rate of 0.0001 to train each network for 5000 epochs. The setup of the task we use is depicted in Figure 5A. This task is challenging because the network must be able to produce different types of sinusoids (i.e., in their frequencies) when presented with inputs that differ only in their constant values.

We trained networks on this task over thirty random network initializations. The trained networks reasonably approximate the sinusoids corresponding to each input (Figure 5C) and were shown to have converged on a solution over the training period (Figure 5D). We found that all types of GLIFR networks performed significantly better than the RNN (Figure 5E). Moreover, learning heterogeneity over training improved model performance both when initialized with heterogeneous parameters (FHetA vs RHetA; two-sample t-test, *p <* 0.001) and when initialized with homogeneous parameters (LHetA vs HomA; two-sample t-test, *p <* 0.001). This suggests that learned heterogeneity improves model performance. This trend holds in models without after-spike currents that are initialized homogeneously (FHet vs RHet, two-sample t-test, *p >* 0.01; LHet vs Hom, two-sample t-test, *p <* 0.001). Interestingly, we found that networks initialized with heterogeneous parameters performed worse than those initialized with homogeneous parameters (RHetA vs LHetA, *p <* 0.001; FHetA vs HomA, *p <* 0.001). This could be because it is difficult to learn the optimal distribution of parameters from an already diverse parameter distribution. However, we observed that the addition of after-spike currents did not produce a statistically significant performance improvement on any types of networks tested (two-sample t-test, *p >* 0.1). Together, these results suggest a computational advantage of biologically realistic dynamics if they are complemented by learning heterogeneous parameters across the network. Interestingly, the addition of after-spike currents was not needed for the improvement over RNNs, as GLIFR networks with solely simpler biological dynamics (e.g., membrane dynamics, voltage reset) performed better than RNNs on this task. While they outperform the RNNs, at least with the simple gradient-based learning used here, the GLIFR networks do not perform as well as the LSTM networks (two-sample t-test, *p <* 0.001).

## 8 Performance on Sequential MNIST (SMNIST) Task

We next assess the performance of the GLIFR network on a Sequential line-by-line MNIST (SMNIST) task. Each image in MNIST is 28 *×* 28. In this task (Figure 6A), the network takes in a 28-dimensional input at each timestep corresponding to a row of the image. At each of the 28 timesteps, the network produces a 10-dimensional output (1 dimension for each digit 0-9), whose softmax would represent a one-hot encoding of the digit. The network is trained so that the 10-dimensional output at the last timepoint correctly indicates the input digit. We use an Adam optimizer with cross entropy loss to train each network for 50 epochs with a learning rate of 0.001. The number of neurons and the number of learnable parameters in each network type trained on this task are shown in Figure 6B.

**Figure 6:**
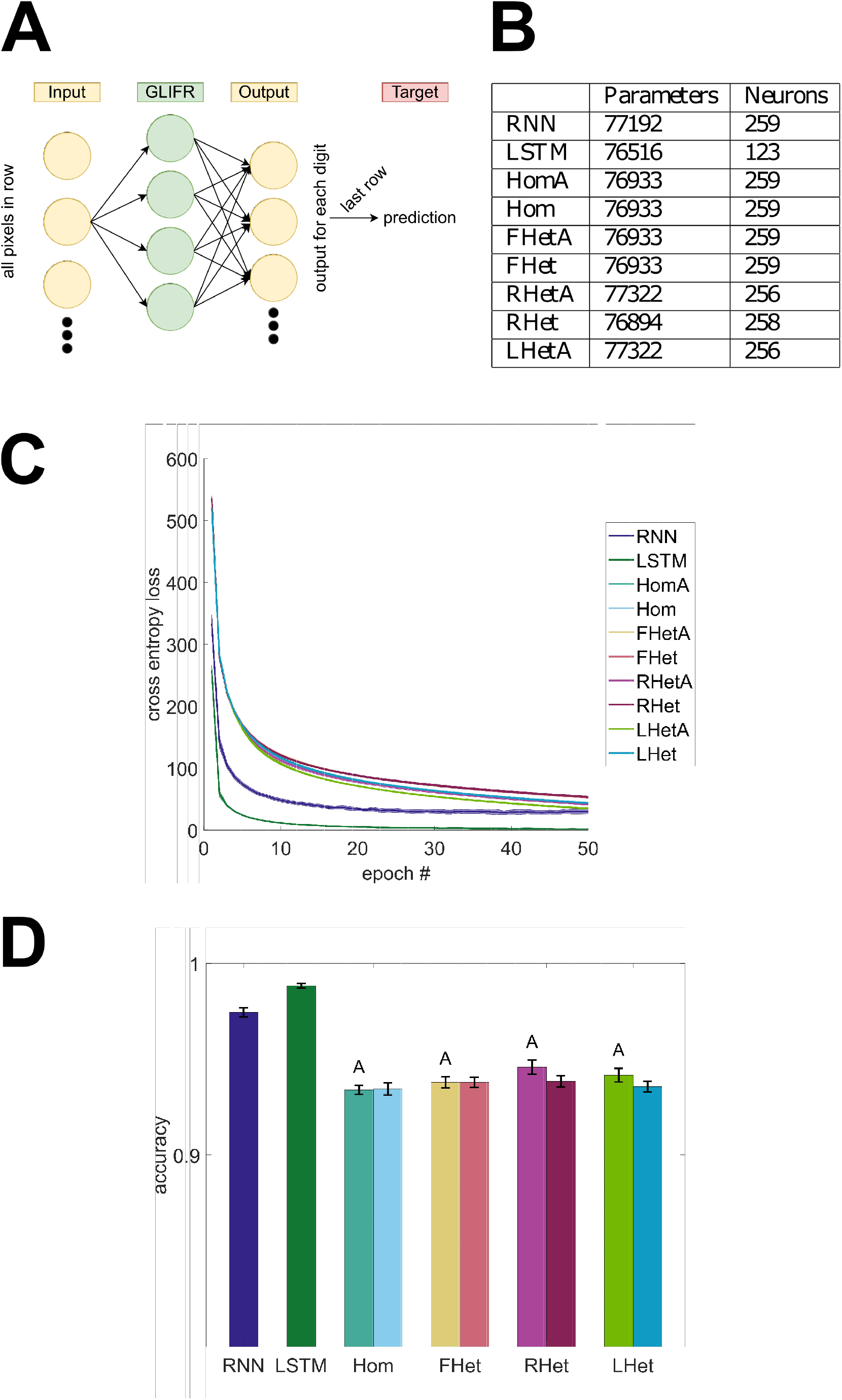
A. SMNIST task. A schematic shows the SMNIST task. The yellow circles represent signals that are either weighted to form inputs to neurons or result from applying weights to neuronal outputs. The green circles represent neurons. The network is input lines of an image at a time and expected to produce, as its output, a one hot encoding of the digit represented in the image. B. Metadata for trained networks. The number of trainable parameters is shown for each network along with the size of the hidden layer. C. Training loss. The training cross-entropy loss of the network averaged over thirty random initializations is plotted over training epochs. The shading indicates a moving average of the standard deviation. On average, all network types converge on a solution within the training time. D. Trained model performance. The accuracy of each trained network on the pattern generation task is shown. Bars marked with “A” above refers to networks that express after-spike currents. Both the RNN and the LSTM networks outperformed the GLIFR networks. However, the ability to learn heterogeneity in the GLIFR networks and the presence of after-spike currents conferred performance improvements.

All network types converged to a solution over the training period (Figure 6C), and we recorded the accuracies of the trained models over thirty random network initializations (Figure 6D). We found that the RNN outperformed all GLIFR networks (two-sample t-test, *p <* 0.001), and that the LSTM outperformed the RNN (two-sample t-test, *p <* 0.001). Similar to the pattern generation task, we found that learning heterogeneity improved performance (FHetA vs RhetA, two-sample t-test, *p <* 0.001; HomA vs LHetA, two-sample t-test, *p <* 0.001) primarily when after-spike currents were included (FHet vs Rhet, two-sample t-test, *p >* 0.1; Hom vs LHet, two-sample t-test, *p >* 0.1). Contrary to our findings in the pattern generation task, networks initialized with heterogeneity performed better than networks initialized without heterogeneity (RHetA vs LHetA, two-sample t-test, *p <* 0.001; FHetA vs HomA, two-sample t-test, *p <* 0.001). However, we found that after-spike currents significantly improved performance in both the RHet (two-sample t-test, *p <* 0.001) and the LHet (two-sample t-test, *p <* 0.001) networks. This suggests that although GLIFR networks achieved lower performance than vanilla RNNs, their complex and diverse dynamics may nevertheless confer a computational advantage.

We further tested the robustness of networks to random silencing of neurons. To accomplish this, for various probabilities p, we performed dropout with probability p during training of each network and tested its performance when a random set of neurons (proportion p) were silenced through the entirety of each testing simulation. While this impaired performance, the networks with GLIFR units are more robust to silencing than vanilla RNNs for *p ≥* 0.4 (Figure 7A), as indicated by thirty random silencing simulations over six random network initializations. The LSTM networks continue to surpass both the GLIFR networks and the RNNs during silencing.

**Figure 7:**
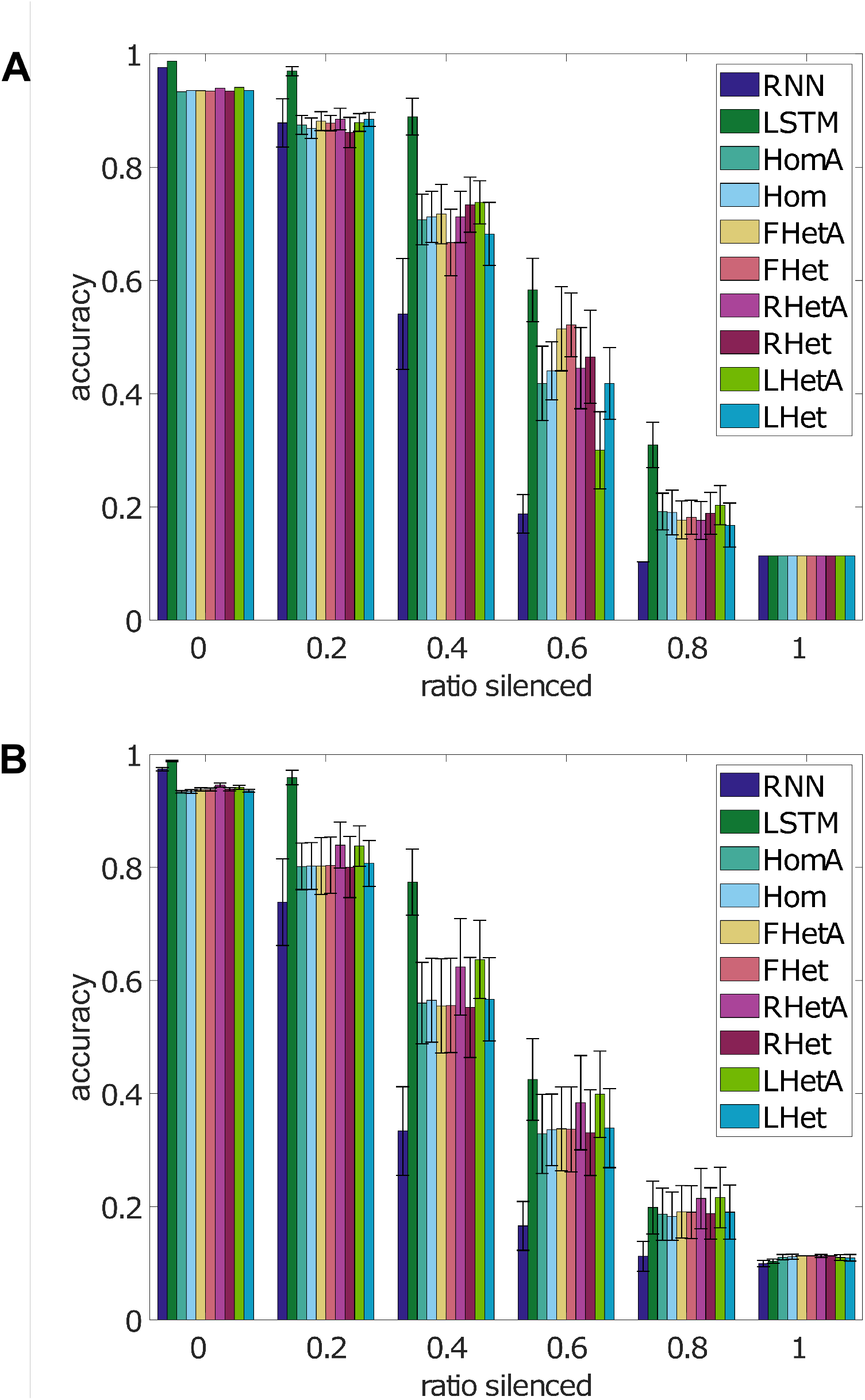
A. Model performance when neurons silenced through training. On five random network initializations, neurons were randomly silenced during SMNIST training for the entire period of a given simulation. Each network was then tested on thirty random silencing of neurons during testing. The average accuracy of each network with varying percentages of their neurons silenced is shown. Bars represent standard deviation. B. Model performance when neurons silenced through testing only. Thirty random network initialization were trained on the SMNIST task. Each network was then tested on thirty random silencing of neurons during testing. The average accuracy of each network with varying percentages of their neurons silenced is shown. Bars represent standard deviation.

GLIFR dynamics enable more complex temporal interactions within and among neurons. What is the resulting impact on the robustness of GLIFR networks random “deletions” (or silencing) of individual neurons? We test this by conducting a similar experiment without dropout through training. Specifically, on networks trained without dropout, we randomly select a subset of neurons to silence throughout testing. For each subset, we clamp the neurons’ firing rates to 0, preventing their contribution in both forward and lateral connections, and compute the accuracy of the network on the MNIST task (Figure 7B). We find that when silencing proportions of neurons for *p ≥* 0.4, all forms of GLIFR networks show an advantage over the RNN. Moreover, in general, LHetA networks perform the best. This observation suggests that neuronal complexity and heterogeneity improves network robustness despite the reduced baseline performance when compared with vanilla RNNs.

## 9 GLIFR Networks Can Learn Discrete Spiking

In all the experiments so far, we have kept *σ*_*V*_ a constant during training. As previously noted, the parameter *σ*_*V*_ can be modified such that as *σ*_*V*_ *<<* 1 the model approaches discrete spiking dynamics. This is established in Equations 14, 15, 16.

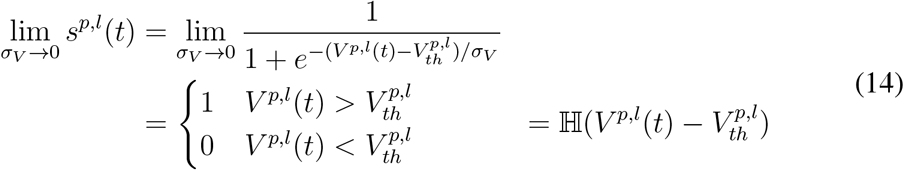

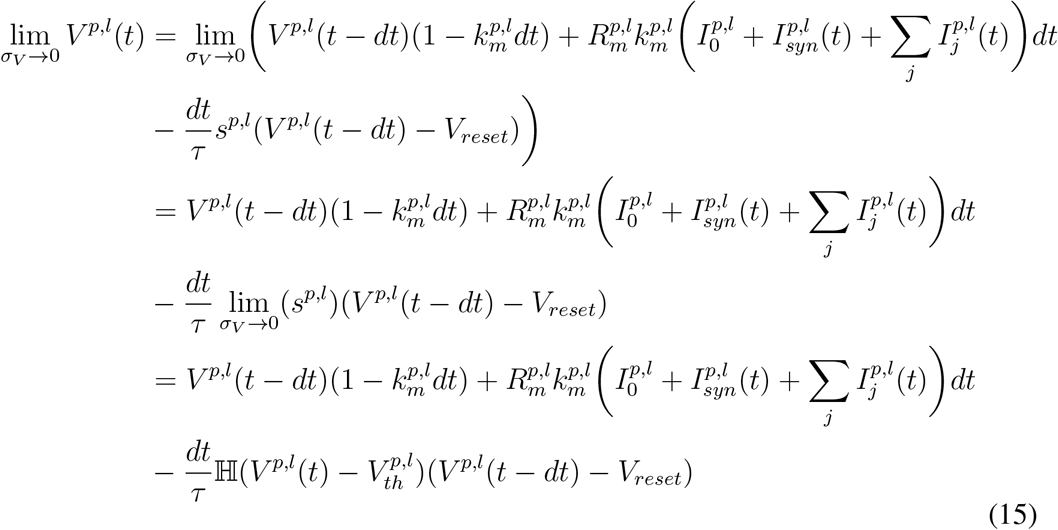

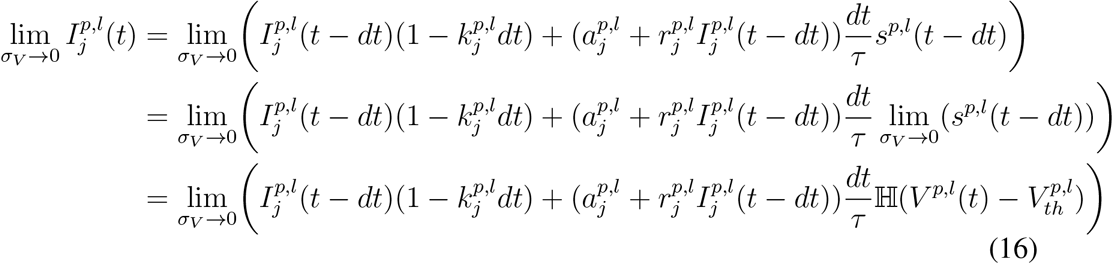

We take a simulated annealing approach to assess whether this setup can be utilized to learn a nearly discrete model which would more closely reproduce biologically realistic, rapid spiking. Specifically, we train a GLIFR network on the SMNIST task while gradually decreasing the *σ*_*V*_ parameter over training (from 1 to 10^−3^). We find that the LHetA networks still learn the task well, achieving an average accuracy of 82.00% (n=30; standard deviation of 0.94%). While this is lower than the previously achieved accuracy of 94.16% without simulated annealing, this annealing experiment demonstrates the ability of GLIFR networks to learn spiking behavior and still perform well without needing to use surrogate gradients.

## 10 GLIFR Networks Learn Heterogeneity Over Training

Based on the performance values above, neuronal heterogeneity seemed to contribute to the GLIFR networks’ performance in the pattern generation task and the SMNIST task. To further elucidate whether and how the learned heterogeneity in parameters may have reshaped neuronal dynamics we determined the extent to which the GLIFR networks had learned truly diverse parameters in both task contexts. For the homogeneously initialized networks, both *a*_*j*_s and *r*_*j*_s had been initialized with limited diversity, and the remaining neuronal parameters had been set homogeneously. We hypothesized that the diversity in all parameters would have developed over training. The heterogeneously initialized networks were set with random distributions of parameters, and we expected the distribution to shift over the course of training.

We found that training did result in heterogeneous parameter values for both the pattern generation task (Figure 8A-B) and the SMNIST task (Figure 8C-D) such that the variance of the trained *a*_*j*_ and *r*_*j*_ parameters is much larger than at initialization (standard deviation was 0.01). We wanted to determine how well this diversity in neuronal parameters mapped to diversity in neuronal dynamics. To do this, we constructed f-I curves representing the average firing rate of each neuron over a time period (5 ms) when the neuron was injected with varying levels of current (Figure 9A, B). We found a diversity in shapes of these curves, illustrating the diversity in the neuronal dynamics produced by neurons in the trained networks.

**Figure 8:**
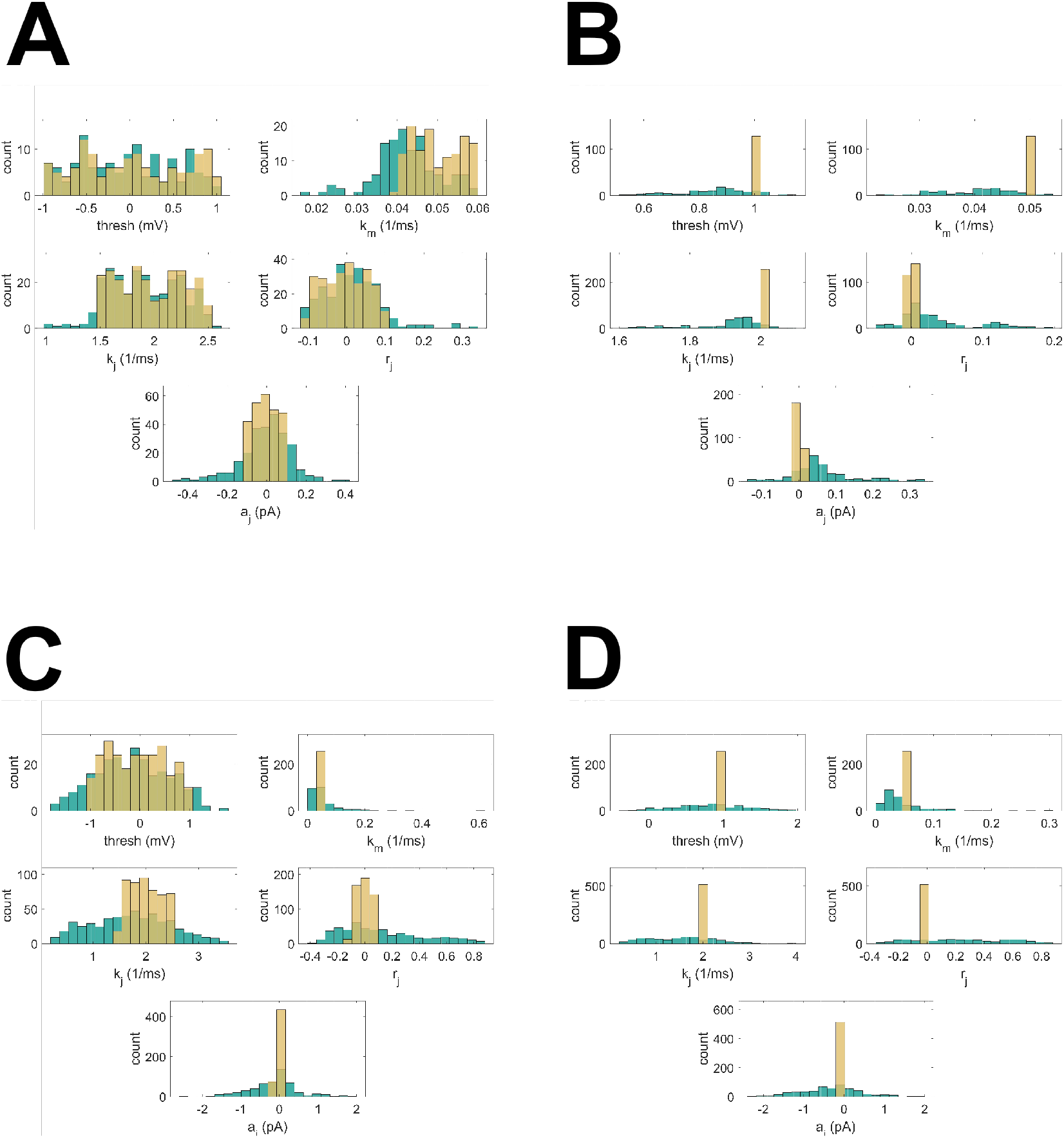
A-D. Learned parameter distributions. The initial (yellow) and learned (green) distributions of parameters across neurons in single example networks are shown for the pattern generation task (A, B) and the SMNIST tasks (C, D). Results are shown for both heterogeneous initialization (A, C) and homogeneous initialization (B, D). Under both initialization schemes, heterogeneity in neuronal parameters is learned.

**Figure 9:**
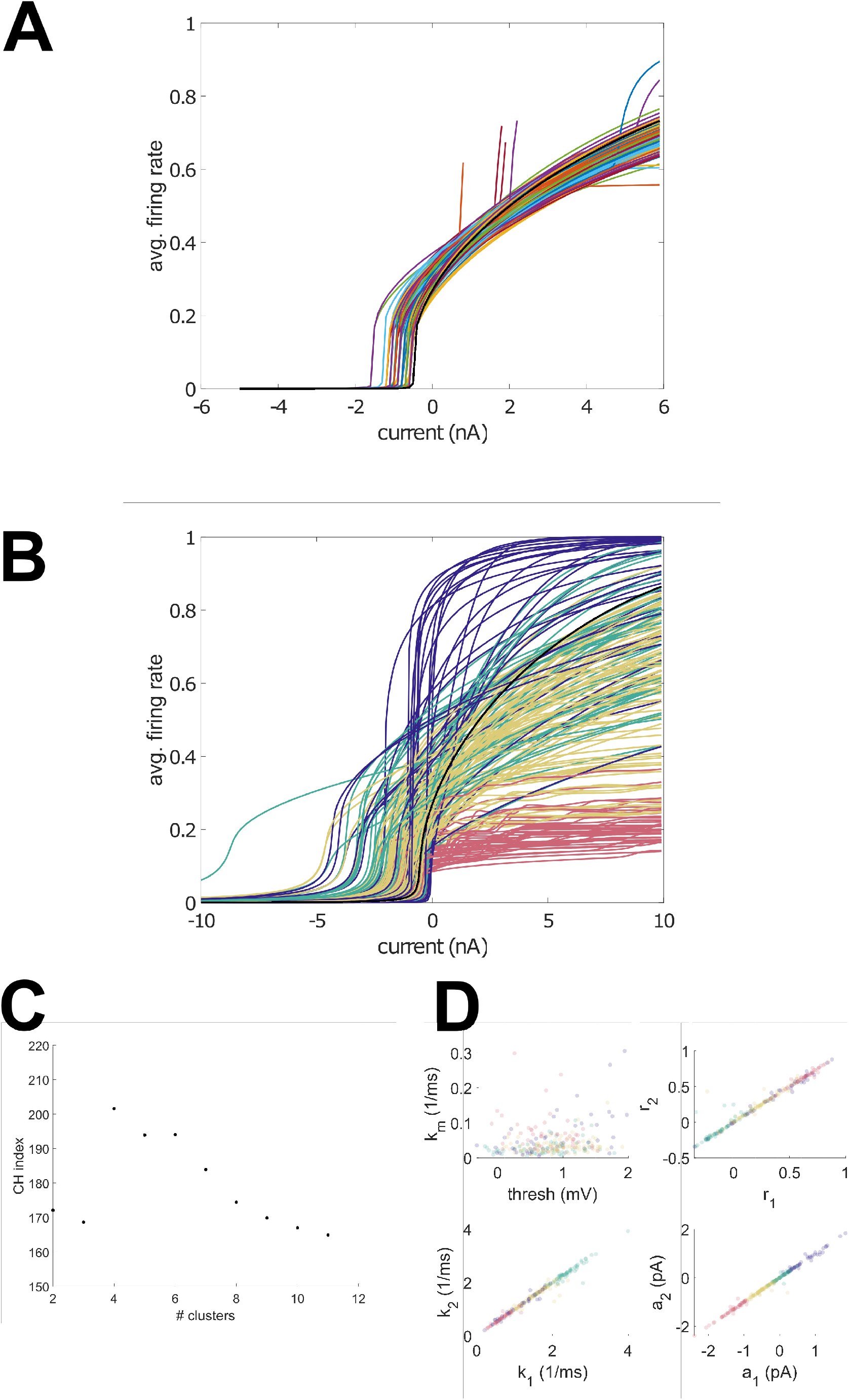
A. Pattern generation task learned dynamics. The f-I curves are plotted for each neuron in an example LHetA network trained on the pattern generation task. The black traces represent those in the initial network. B. SMNIST task learned dynamics. The f-I curves are plotted for each neuron in the LHetA network trained on the MNIST task and are colored according to clusters produced by Ward’s hierarchical clustering (colors correspond with those in D) This demonstrates that the clustering of the neuronal parameters also separates the f-I curves that characterize the neuronal dynamics. C. CH scores. The CH scores for different cluster numbers are plotted for SMNIST parameters, demonstrating an optimal cluster count of 4. D. Parameter clustering. Various pairs of parameters are plotted for the LHetA network trained on the SMNIST task. The points are colored according to clusters produced by Ward’s hierarchical clustering.

Finally, we hypothesized that networks would also develop classes of neurons. To that end, we performed hierarchical clustering on the neuronal parameters of networks trained on the SMNIST task. We used the Calinski-Harabasz (CH) Index as a measure of clustering. For the pattern generation task, we were unable to find a number of clusters that produced the optimal CH score. However, we found that the CH score was maximized using four clusters in the LHet network trained on the MNIST task (Figure 9C). After clustering these neurons, we examined their parameters. The four classes of neurons appeared to be separated primarily based on after-spike current parameters *a*_*j*_ and *k*_*j*_ (Figure 9D). Specifically, one class expresses slow, after-spike currents with hyperpolarizing values of *a*_*j*_ and depolarizing values of *r*_*j*_ (red), a second class expresses smaller hyperpolarizing after-spike currents (yellow), a third class expresses slow depolarizing after-spike currents (purple), and the final class expresses fast after-spike currents with depolarizing values of *a*_*j*_ and hyperpolarizing values of *r*_*j*_ (green). We noted that each pair of parameters corresponding to after-spike currents tended to be similar (i.e., for a given neuron, *a*_1_*≈ a*_2_). We suggest that this is a result of the gradient descent technique. Potentially, the gradients over both after-spike currents were similar, resulting in both sets of parameters progressing in the same direction.

We also found that the classes based on parameters separated the f-I curves into categories (Figure 9B). For example, the neurons with hyperpolarizing after-spike currents tended to exhibit low maximal firing rates. On the other hand, the neurons with slow depolarizing after-spike currents tended to exhibit firing rates that rapidly saturated to relatively high values. This further suggests that the learned diversity in parameters enabled a diversity in dynamics as well.

## 11 Discussion

This work explored the role of neuronal complexity and heterogeneity in network learning and computation using a novel paradigm. Specifically, we developed the GLIFR model, a differentiable rate-based neuronal model that expresses after-spike currents in addition to the simple types of dynamics. While past work (Perez-Nieves et al., 2021; Salaj et al., 2021) has studied neuronal complexity and heterogeneity, it is beneficial to learn dynamics in neurons using traditional machine learning approaches. Here we demonstrated the ability to learn both synaptic weights and individual neuronal parameters underlying intrinsic dynamics with traditional gradient descent. While it is generally rate-based, the GLIFR model retains the ability to produce spiking outputs and thus is a powerful model for studying neuronal dynamics.

We demonstrated the ability for networks of GLIFR neurons to learn tasks, such as a pattern generation task and a SMNIST task. In both tasks, heterogeneous parameters and dynamics were learned. We tested the effects of the ability to learn parameter heterogeneity and the ability to express after-spike currents on model performance. Learning heterogeneous parameter distributions improved performance, and modeling after-spike currents improved performance on the SMNIST task. Regardless of whether heterogeneous parameters or after-spike currents were learned, the GLIFR models outperformed vanilla RNNs in the pattern generation task. We also found that GLIFR networks trained on the SMNIST task were more robust to injury. Specifically, when we silenced fixed fractions of neurons, the GLIFR networks performed better than vanilla recurrent neural networks. Finally, we found that learning parameters across a network in response to a temporally challenging task enabled the network to develop classes of neurons with differing intrinsic parameters and dynamics.

These findings support the hypothesis that neuronal heterogeneity and complexity have a computational role in learning complex tasks. The implications of this are two-fold. On one hand, neuronal heterogeneity may be an interesting step to pursue for more powerful computing in artificial neural networks. Vanilla recurrent networks, for example, rely on a single type of dynamic - typically either the sigmoid or the tanh activation function. However, the use of activation functions that can be “learned” over time, such that the trained network exhibits diverse dynamics across its neurons, may confer a computational advantage. Here we demonstrated the computational advantages of more biologically realistic neurons for specific temporally challenging tasks while using traditional gradient descent training techniques.

On the other hand, our results provide further insight into the purpose of complexity and diversity of neural dynamics seen in the brain. Our brains are responsible for integrating various sensory stimuli, each varying in their temporal structures. Intuitively, heterogeneity in the types of dynamics used by neurons across the brain may enable robust encoding of a broad range of stimuli. Moreover, certain types of complex dynamics may affect synaptic strengths and improve the network’s robustness to damage. Along these lines, our results suggest that neuronal complexity and heterogeneity play a role in network function.

We believe that with the GLIFR model in public domain, we have opened door to more intriguing studies that will identify further roles for complex neural dynamics in learning tasks. In future work, testing the GLIFR models on additional tasks may provide additional insights into the computational advantages of neuronal heterogeneity and complexity. Additionally, it would be interesting to pursue a theoretical explanation of the computational role of heterogeneity and complexity. Past experimental work (Tripathy, Padmanabhan, Gerkin, & Urban, 2013) found that neuronal heterogeneity increased the amount of information carried by each neuron and reduced redundancy in the network. Exploring this in computational models would be valuable, as it may suggest additional ways in which the computational advantages of biological dynamics can be harnessed to improve artificial neural networks and yield insights into the mechanisms of computation in biological networks.

## 12 Code availability

Modeling, training, and analysis code is available at https://github.com/AllenInstitute/GLIFS_ASC.

## 13 Acknowledgments

We wish the acknowledge the following sources of funding and support: the University of Washington Mary Gates Research Endowment, the NIH Training Grant T90 DA 32436-10, the NIH BRAIN Initiative Grant 1RF1DA055669. We wish to thank the Allen Institute founder, Paul G. Allen, for his vision, encouragement, and support.

